# Formaldehyde induces and promotes Alzheimer’s disease pathologies in a 3D human neural cell culture system

**DOI:** 10.1101/2025.02.27.640690

**Authors:** Peipei Wu, Danqi Chen, Fei Wang, Kun Lu, Einar M Sigurdsson, Chunyuan Jin

**Affiliations:** Department of Medicine, New York University Grossman School of Medicine, New York, NY 10016, United States; School of Public Health, China Medical University, Shenyang 110122, China; Department of Environmental Sciences and Engineering, Gillings School of Global Public Health, University of North Carolina, Chapel Hill, NC, 27599, United States; Department of Neuroscience and Physiology, Neuroscience Institute, New York University Grossman School of Medicine, New York, NY 10016; Department of Psychiatry, New York University Grossman School of Medicine, New York, NY 10016, United States

## Abstract

Alzheimer’s disease (AD) arises from complex multilevel interactions between genetic, epigenetic, and environmental factors. Recent studies suggest that exposure to the environmental and occupational toxicant formaldehyde (FA) may play a significant role in AD development. However, the effects of FA exposure on Aβ and tau pathologies in human neural cell 3D culture systems remain unexplored. To investigate FA’s role in AD initiation, we differentiated 3D-cultured immortalized human neural progenitor ReN cells (ReNcell VM) into neurons and glial cells, followed by FA treatment. FA exposure for 12 weeks resulted in a dose-dependent increase in Aβ40, Aβ42, and phosphorylated tau levels. To further examine FA’s role in AD progression, we established a 3D human neural cell culture AD model by transfecting ReN cells with AD-related mutant genes, including mutant APP and PSEN1, which recapitulate key AD pathological events. Our findings demonstrate that FA exposure significantly elevated Aβ40, Aβ42, and phosphorylated tau levels in this 3D-cultured AD model. These results suggest that FA exposure contributes to the initiation and progression of AD pathology in 3D-cultured human neural cells.

## Introduction

Alzheimer’s disease (AD), a chronic neurodegenerative disorder characterized by progressive executive and cognitive dysfunction as well as behavior impairment, accounts for 60 to 70 percent of dementia cases (Masters et al., 2015; Scheltens et al., 2016; Wilson et al., 2012). Accumulation of amyloid-β (Aβ) plaques and intracellular neurofibrillary tangles (NFTs) of hyperphosphorylated tau protein (p-tau) are two main hallmarks of AD. Due to the complexity of human brains, and limitations of animal models and research tools, the detailed pathogenesis of AD remains incompletely understood. Many hypotheses about AD have been proposed, including those involving Aβ accumulation, tau pathology, cholinergic neuron damage, oxidative stress, and inflammation. However, most preventive or therapeutic strategies targeting various pathologies in AD remain relatively ineffective, which has led to an urgent need to investigate its etiology from a different angle.

Multilevel complex interactions between genetic, epigenetic, and environmental factors contribute to the occurrence and progression of AD. Formaldehyde (FA), the simplest saturated aldehyde with protein cross-linking properties, is one of the most common environmental and occupational aldehyde pollutants. FA is widely present in the environment and can be found in construction materials, agricultural fertilizers, fumigants, paints, cosmetics, polishes, and cleaning agents (Tong, Han, Luo, Li, et al., 2013). Importantly, FA is found to gradually accumulate in brains during the aging process (Tong, Han, Luo, Wang, et al., 2013). It has been shown to be associated with AD and dementia in growing numbers of epidemiological, animal, and *in vitro* studies. A recent large cohort study of over 75,000 participants, including more than 6,000 FA-exposed workers, demonstrated an increased risk of cognitive impairment in FA-exposed individuals (Letellier et al., 2022). Another epidemiological study of 305 women working as histology technicians, who were chronically exposed to FA, demonstrated a marked decline in cognitive function (Kilburn et al., 1987). Several other small-scale studies also demonstrated a link between FA exposure and impaired memory and intellectual performance (Kilburn et al., 1987; Kilburn & Warshaw, 1992; LoSasso et al., 2001). Moreover, elevated levels of uric FA have been shown to correlate positively with the severity of cognitive impairment in AD patients (Tong et al., 2011). FA concentrations in the hippocampus of AD patients are significantly higher than those in healthy aged individuals, suggesting its potential as a biomarker for AD (Tong et al., 2017). Additionally, endogenous FA accumulation has been shown to impair memory in patients with mutations in the FA-metabolizing enzyme ALDH2, as well as in ALDH2 knockout mice (Ai et al., 2019).

Animal experiments have implicated causative role of FA exposure in AD development and progression. In young rhesus monkeys, excess FA induced by methanol feeding led to memory decline, increased tau protein phosphorylation at residues T181 and S396 in cerebrospinal fluid, and the formation of p-tau aggregates and Aβ plaques in the brain (Yang et al., 2014). In rhesus monkeys, chronic exposure to FA was found to induce major AD-like pathological markers and cognitive impairments, including the presence of Aβ and neuritic-like plaques, NFTs, increased p-tau, neuronal loss, and reactive gliosis in three brain regions associated with memory and AD (Zhai et al., 2018). Rodent studies have also shown that acute FA exposure induces early AD-like changes, including cognitive deficits, pathological alterations in the brain, accumulation of Aβ, and p-tau in the cerebral cortex (Liu et al., 2018). Moreover, FA injection has been shown to suppress hippocampal long-term potentiation (LTP) by blocking N-methyl-D-aspartate (NMDA) receptors, leading to impaired spatial memory performance in normal Sprague-Dawley rats (Tong, Han, Luo, Wang, et al., 2013). While these animal experiments strongly suggest that FA exposure is critical for AD pathology, its role in human AD pathogenesis has yet to be thoroughly examined.

By transducing both APP and PSEN mutants, Choi et al. successfully recapitulated AD pathologies, including robust Aβ deposition and NFT-like tau pathology, in a 3D human neural cell culture system (Choi et al., 2014). This system may more accurately recapitulate *in vivo* human AD pathology than transgenic mouse models or 2D human cell models of AD, providing a valuable platform for mechanistic studies in a human-brain-like environment. In this study, we demonstrate that FA exposure induces AD pathologies in the 3D human neural cell culture system, highlighting the potential role of FA in the initiation and progression of AD.

## Methods

### Cell culture and treatment

The human neural progenitor cell line (ReNcell VM) was purchased from EMD Millipore. ReNcell 2D or 3D culture and transfection were performed following an established protocol (Kim et al., 2015). For the 2D culture, ReN cells were seeded onto Matrigel precoated plates and cultured in DMEM/F12 medium (Gibco/Life Technologies) supplemented with Heparin (Stemcell Technology), B27 (Gibco/Life Technologies), bFGF (REPROCELL Inc.), and EGF (Millipore Sigma) at 37°C and 5% CO_2_. Differentiation was initiated by removing growth factor from the medium once the cells reached 70% confluency. After two weeks, differentiated ReN cells (2D-culture) were treated with 0 and 100 μM FA for 7 days.

To establish the 3D cultured AD model, we obtained the CMV-APP-GFP, which carries both K670N/M671L (Swedish) and V717I (London) FAD mutations (APPSL), and the CMV-PSEN-RFP vector, which contains the ΔE9 FAD mutation (PS1ΔE9) (Millipore). The lentiviral polycistronic vectors CSCW-IRES-GFP and CSCW-IRES-mCherry were purchased from MGH Vector Core Facility. The cells were transfected with CMV-APP-GFP and CMV-PSEN-RFP together. ReN cells with CSCW-IRES-GFP or CSCW-IRES-mCherry alone were also generated as controls. Viral infection and enrichment of high-expressing transgenic ReN cells were performed according to a protocol for lentivirus infection (EMD Millipore). Cells were sorted using a 100 μm nozzle with a sheath pressure of 20 psi. Parameter settings: GFP-488 nm, mCherry-561 nm. Gates were set to select the top 2% of high-expressing cells. The enriched transgenic ReN cells were plated on a thick-layer within 6-well plate inserts (Falcon® Cell Culture Inserts, Corning, NY) and on a thin-layer within 4-well culture slides (Falcon® 4-well Culture Slide, Corning, NY). The groups were divided as follows: ReN (control), ReN cells treated with 100 μM (ReN FA100), ReN cells treated with 200 μM FA (ReN FA200), ReN cells transfected with GFP and mCherry plasmids (G+M), ReN cells transfected with GFP and mCherry plasmids and treated with 50 μM FA (G+M FA50), 100 μM FA (G+M FA100), and 150 μM FA (G+M FA150). Additional groups included ReN cells transfected with mutant APP and PSEN (AD), ReN cells transfected with mutants APP and PSEN and treated with 0 μM FA (AD FA0), 100 μM FA (AD FA100), and 150 μM FA (AD FA150). The treatment lasted for up to 4 months.

### Animal exposure

Rat exposure was performed at the Lovelace Respiratory Research Institute (Albuquerque, NM, USA) according to approved protocols for the use of vertebrate animals in experiments. Animal study was conducted in accordance with the National Institutes of Health guidelines for the care and use of laboratory animals (Leng et al., 2019). Fischer 344 rats (6-week-old, male) were exposed to isotope-labeled [^13^CD_2_]-FA vapor by single-exposure, nose-only inhalation with the final target exposure concentration of 0 (air control) or 2 ppm for 28 days (6 h/day). The concentration of the exposure chamber was monitored by collection of Waters XpoSure Aldehyde Sampler cartridges every 10 min continuously throughout the exposure. Animals were euthanized using a pentobarbital-based euthanasia solution to induce surgical level anesthesia, followed by the introduction of pneumothorax. Brain tissue samples were then harvested, wrapped in aluminum foil, and immediately snap-frozen in liquid nitrogen. Samples were stored at −80 °C until further analysis.

### Cell proliferation assay

The 3-(4,5-dimethylthiazol-2-ul)-2,5-diphenyl tetrasodium bromide (MTT, Sigma-Aldrich) cytotoxicity assay measures mitochondrial reductases, which converts the water soluble MTT salt to a formazan that accumulates in healthy cells. ReN cells in proliferation state were digested using accutase, an enzyme mixture used for detaching adherent cells (Gibco/Life Technologies). A total of 2500 cells per well were seeded into a 96-well plate and treated with 0, 25, 50, 100, and 200 μM FA for 2, 4, 6, and 8 days. Medium was changed every 2 days. At the end of each timepoint, 10 µl MTT was added to each well containing 100 µl fresh medium with final concentration of 5 mg/ml MTT and incubated for 4 h at 37°C in 5% CO_2_. At the end of the incubation, 100 µl of isopropanol containing 0.04 N HCl was added to each well to dissolve the crystals. The absorbance of wells was read at 570 nm using a microplate reader.

### ELISA

Soluble Aβ40 and Aβ42 levels in the conditioned medium were measured using Human/Rat amyloid-β ELISA Kit from Wako (Osaka, Japan). Media from undifferentiated and differentiated ReN cells were collected and diluted 1:2 with the dilution buffer provided by the kit. Absorbance was measured at 450 nm using a microplate reader.

### Western blot analysis

The 2D-cultured ReN cells were lysed using 150 μL of a boiling buffer containing 0.5 mM dithiothreitol (DTT), protease inhibitors (PMSF 100X, Aprotinin 1000X, Leupeptin 1000X; all from Sigma) and phosphatase inhibitor cocktail 100X (Cell Signaling Technology), followed by sonication to get a homogenized lysate. Protein concentration was quantified using the Pierce™ BCA protein assay kit (Thermo Fisher Scientific, Waltham, MA, USA) according to the manufacturer’s instructions. The 3D-cultured cells were collected and washed with pre-cooled PBS, then homogenized in the boiling buffer containing protease inhibitors and phosphatase inhibitor cocktail and incubated on ice for 10 min. This was followed by sonication for 5 min to ensure thorough lysis.

For rat brain tissue lysate, 0.1 g of brain tissue for each sample was shredded and cleaned using pre-cooled PBS, followed by sonication in a boiling buffer to obtain homogenate lysate. Lysate was then mixed with 6× SDS sample buffer and boiled at 95 °C for 10 min. A total of 40 μg of whole cell lysate or 50 μg of tissue lysate per lane was separated using 12% SDS-PAGE or a 12% gradient gel (Invitrogen, Waltham, MA, USA), respectively. Blots were transferred to a polyvinylidene difluoride membrane, 0.45 μm (Bio-Rad, Hercules, CA, USA), blocked with 5% milk (Blotting-Grade Blocker, Bio-Rad, Hercules, CA, USA) in TBST for 30 min at room temperature, and then incubated in primary antibodies (see Table 1) overnight at 4 °C. After being washed with TBST for 10 min three times, the membranes were incubated with a secondary antibody conjugated with horseradish peroxidase for 1 h at room temperature. Visualization was performed using the ECL Plus Western blotting detection reagent (Bio-Rad, Hercules, CA, USA), and quantification of the bands was performed using NIH Image J software.

**Table 1:** The primary antibodies used for western blot:

| Name | Source | CatLog number | Dilution |
| --- | --- | --- | --- |
| GAPDH | Santa Cruz | sc- 47724 | 1:2000 |
| β-actin | Abcam | AB8226 | 1:2000 |
| p-Tau AT8 | Thermo Scientific | MN1020 | 1:1000 |
| p-Tau Ser199 | Thermo Scientific | 44-734G | 1:1000 |
| Total tau | Santa Cruz | 136400 | 1:2000 |
| APP (Aβeta 6E10) | Covance | SIG-39300 | 1:2000 |

### Statistical analysis

All data were expressed as mean ± standard deviations (SD), significant differences among and between group means were carried out using GraphPad Prism (San Diego, CA). Significant differences between the means were evaluated by analysis of variance test (unpaired t test, 1-way ANOVA or 2-way ANOVA) as indicated in the figure legends. Post hoc tests, when appropriate, were analyzed using Tukey’s multiple comparisons test. Statistical significance was defined as p < 0.05.

## Results

### FA exposure induces AD pathologies in human neural cells

Human neural progenitor ReN cells differentiate into neurons and glial cells in the absence of growth factors in the culture medium (Kim et al., 2015). We first evaluated the cytotoxicity of FA in proliferating ReN cells by treating them with varying doses of FA (0, 25, 50, 100, and 200 μM) for 8 days. Lower doses of FA (0–100 μM) did not elicit significant changes in cell viability, whereas 200 μM caused approximately 30% cell death by day 8 (Fig. 1A). Based on these findings, we focused on the effects of 100 μM FA in 2D culture experiments. This concentration is physiologically relevant, as FA levels in human blood range between 20 μM and 100 μM (Heck et al., 1985; Luo et al., 2001; Nagy et al., 2004). To investigate whether FA induces AD pathologies, we exposed differentiated ReN cells to 100 μM FA for 7 days and then monitored the expression levels of p-tau and APP. Western blot analysis showed that FA exposure increased the levels of p-tau (Ser199) and p-tau (AT8, Ser202&Thr205) compared to the control, while total tau levels remained unchanged (Fig. 1B, C; Fig. S1). Additionally, APP levels increased following FA exposure (Fig. 1D, E; Fig. S2). These results indicate that FA can induce AD-like pathologies in human neural cells.

**Figure 1.**
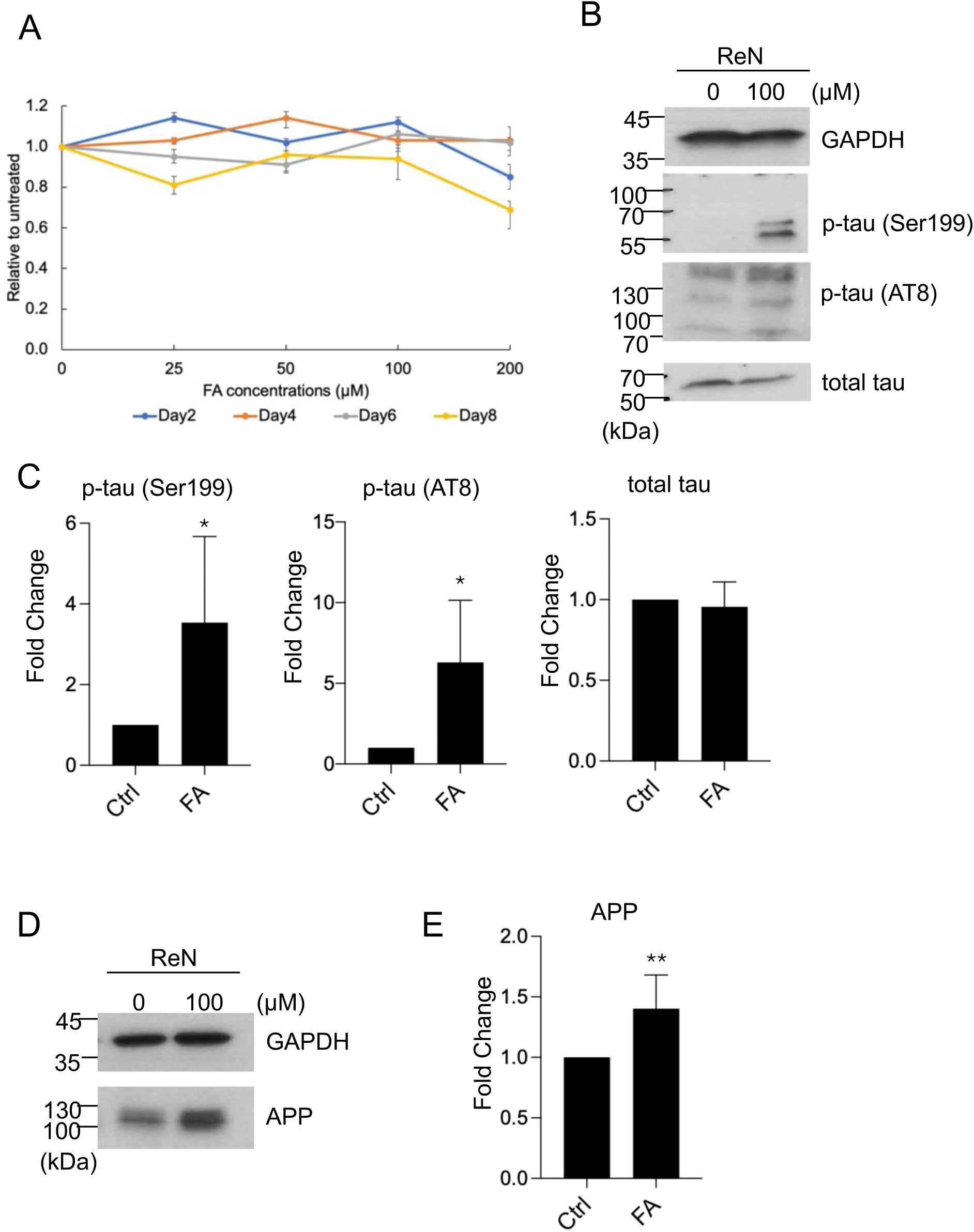
FA induces AD pathologies in human neuronal cells. **(A)** Cytotoxicity of FA in ReN cells was tested using MTT assays. **(B–E)** ReN cells were treated with or without 100 μM FA for 1 week following 2 weeks of differentiation initiation. Whole cell lysates were extracted followed by Western blot to assess tau phosphorylation at different sites, total tau, and APP levels (B, D). The levels of p-tau phosphorylation, total tau, and APP were quantified and normalized to GAPDH, with the relative intensity in the control (Ctrl) set to one (C, E). Unpaired t test was applied to compare differences between Ctrl and FA. Results are presented as mean ± SEM (n = 3). \**p* < 0.05, \*\**p* < 0.01.

### FA exposure induces AD pathologies in rat brain tissues

To investigate if this occurs *in vivo*, male rats were exposed to 2 ppm of FA for 28 days, after which brain tissues were collected for Western blot analysis using antibodies against p-tau and total tau. The results showed that p-tau (Ser199) levels were elevated in FA-exposed rats (n=4) compared to control rats (n=2), while total tau levels remained unchanged (Fig. 2A-D; Figs. S3 and S4). These findings suggest that FA exposure induces tau hyperphosphorylation *in vivo*.

**Figure 2.**
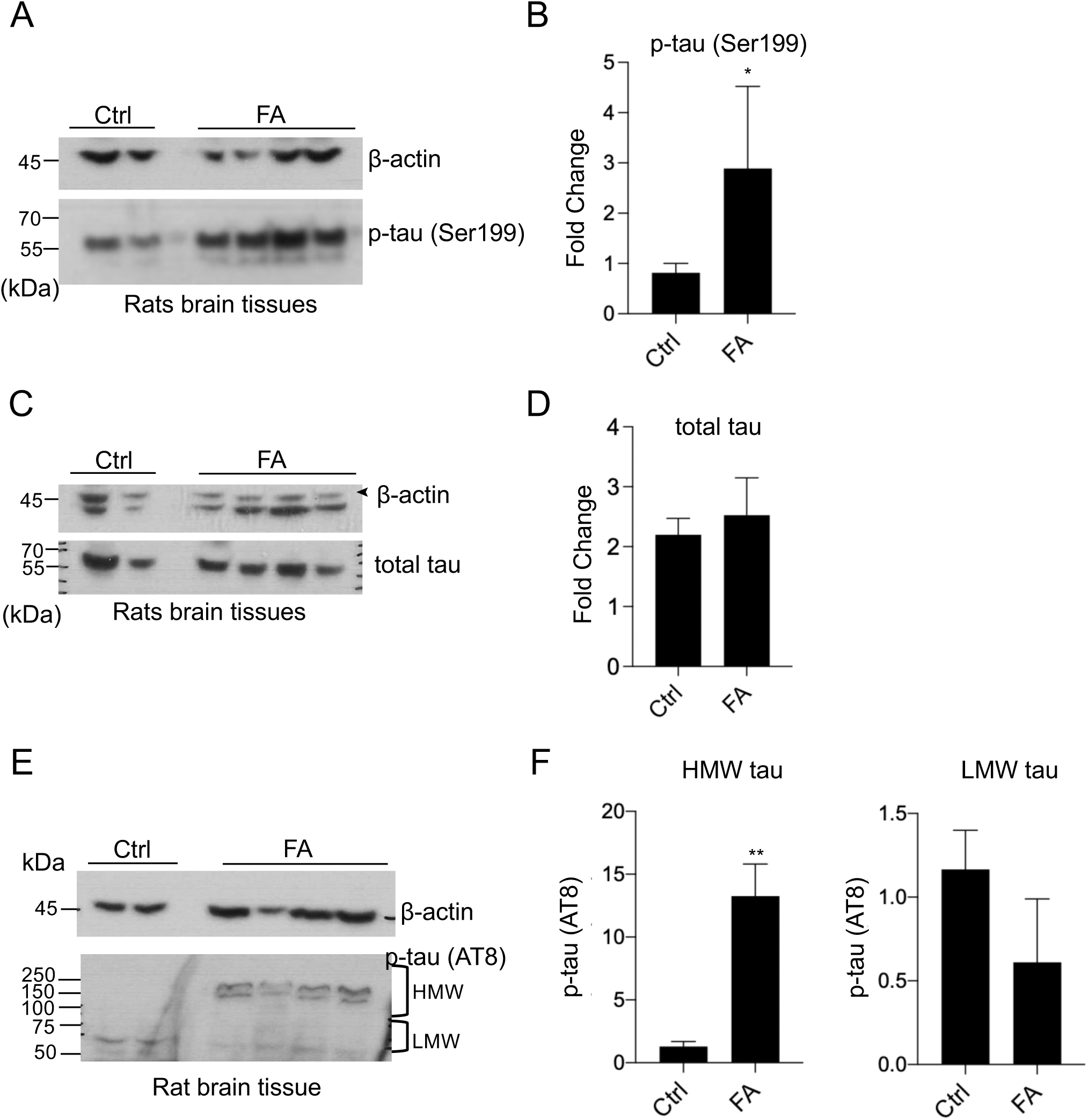
FA exposure induces AD pathologies in rat brain tissues. **(A, C, E)** Rats were exposed to 2 ppm FA for 4 weeks. Whole cell lysates were extracted from brain tissues followed by Western blot to assess p-tau, total tau, and β-actin levels. **(B, D, F)** Quantification of p-tau levels in Ctrl (n = 2) and FA (n = 4) groups showed increased p-tau at Ser199 and HMW-AT8 immunoreactive tau in the FA-exposed group compared to the control group. The levels of p-tau phosphorylation and total tau were quantified and normalized to β-actin, with the relative intensity in the Ctrl set to one. Unpaired t test was applied to compare differences between Ctrl and FA. Results are presented as mean ± SEM (n = 3). \**p* < 0.05, \*\**p* < 0.01.

Tau protein can be divided into high-molecular-weight tau oligomers (HMW-tau) and low-molecular-weight tau (LMW-tau). Recent studies have reported that HMW-tau is significantly elevated in post-mortem AD brains and AD animal models, as observed in western blot analyses (Kopke et al., 1993; Miao et al., 2019; Zhou et al., 2018). Interestingly, the p-tau antibody AT8 recognized multiple bands in our rat brain tissue samples: lower bands (50–75 kDa), likely corresponding to LMW-tau, and higher bands (150–250 kDa), likely corresponding to HMW-tau (Fig. 2E; Fig. S5). Based on our Western blot results, phosphorylation of HMW-tau at Ser202&Thr205, detected by AT8 antibody, was significantly elevated in FA exposed rats, whereas phosphorylation of LMW-tau showed a slight, non-significant decrease (Fig. 2E, F; Fig. S5). Based on previous study, HMW-tau is selectively present in AD brains, and was hyperphosphorylated at multiple sites (Zhou et al., 2018). This evidence supports our findings that FA exposure leads to AD-like tau pathologies in rat brain tissue—tau abnormal hyperphosphorylation and aggregation which forms oligomers and paired helical filaments.

### FA exposure increases AD pathologies in a 3D human neural cell culture system

To better mimic the *in vivo* brain environment, ReN cells were differentiated and cultured in a three-dimensional (3D) matrix by embedding them in Matrigel. Matrigel can provide cells with multiple structural proteins and a brain-like stereo structure, which can accelerate neuronal differentiation and neural network formation (Hughes et al., 2010; Ortinau et al., 2010; Tang-Schomer et al., 2014). To investigate the effect of FA on AD initiation, we first induced differentiation of ReN cells by removing growth factors from the proliferation medium for three weeks. The differentiated ReN cells were then continuously cultured in the 3D environment in the presence of 0, 100, and 150 μM FA for 6 to 12 weeks. These doses used in 3D culture are considered physiologically relevant, as the FA concentration in the brain is approximately 200 μM in the hippocampus and 400 μM in the cortex of healthy individuals (Tong, Han, Luo, Wang, et al., 2013). The levels of soluble Aβ40 and Aβ42 in the conditioned media were measured by ELISA. ReN cells treated with 100 and 150 μM FA showed significant increases in Aβ40 and Aβ42 as compared to the control at 6 and 12 weeks (Fig 3A, B). Both Aβ40 and Aβ42 increased in a dose-dependent manner at 12 weeks (Fig. 3A, B). Western blot analysis was then conducted to detect AD markers in ReN cells exposed to 0, 100, and 150 μM FA for 12 weeks. The levels of phosphorylated tau at Ser199 and Ser202&Thr205 dramatically increased following FA exposure in a dose-dependent manner (Fig. 3C-F; Figs. S6 and S7). Moreover, the levels of APP, the primary source of Aβ peptide, were also elevated following FA exposure compared to control cells (Fig. 3G, H; Fig. S8). These results indicate that FA exposure induces AD pathologies in

**Figure 3.**
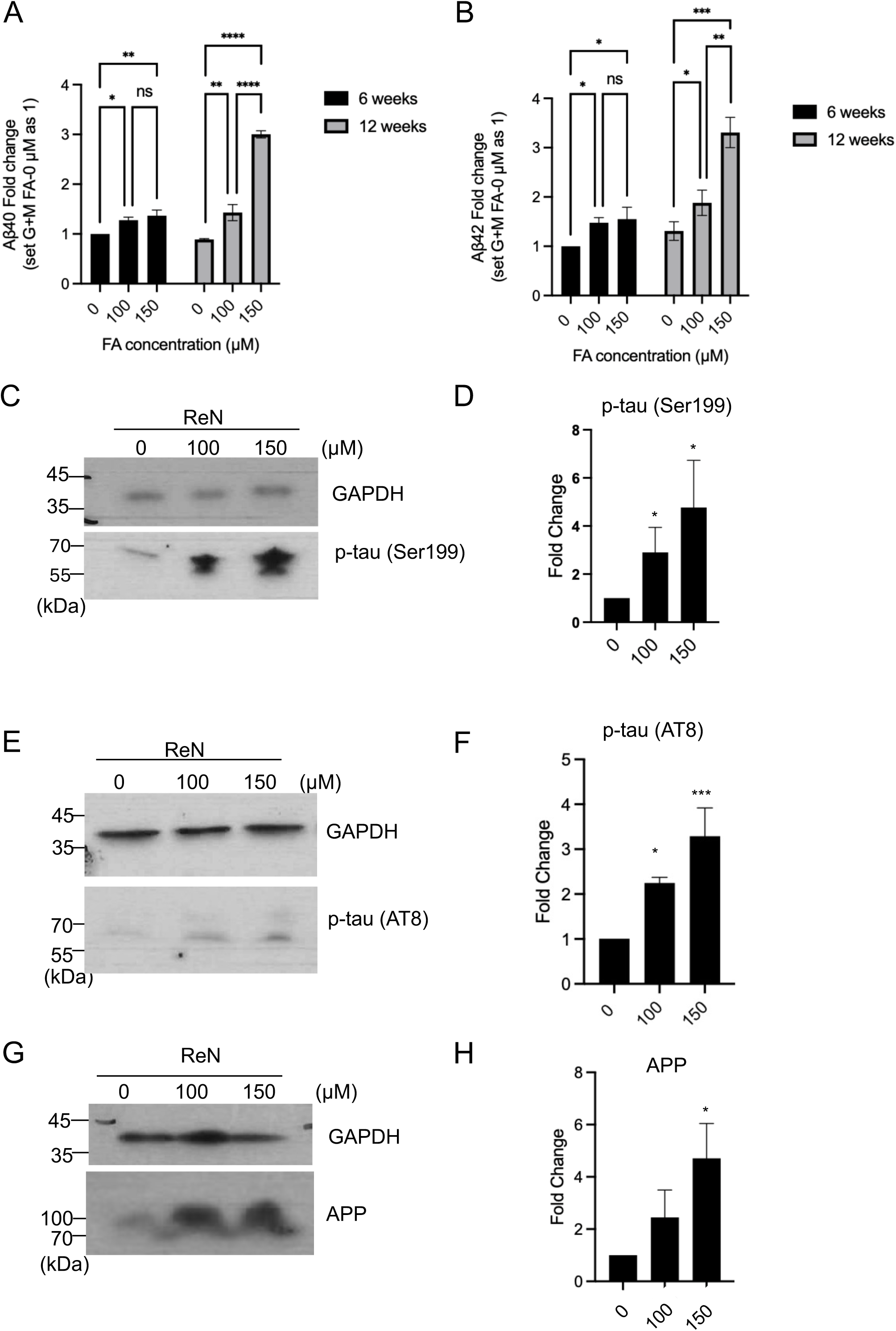
Long-term FA exposure increases AD pathological makers in 3D-culture ReN cells. **(A, B)** After differentiation, 3D-cultured ReN cells were treated with 0, 100, 150 μM FA. Amyloid-β peptides (Aβ40 and Aβ42) in the conditioned medium were measured by ELISA after 6 or 12 weeks of FA exposure and normalized to protein concentration. Soluble Aβ40 **(A)** and Aβ42 **(B)** levels were increased by FA after 6 weeks of exposure, and both Aβ40 and Aβ42 showed a dose-dependent increase at 12 weeks of exposure. One-way analysis of variance (ANOVA) followed by Turkey’s multiple comparisons test was applied to compare differences between doses at 6 weeks and 12 weeks separately. Results are presented as mean ± SEM (n = 3). \**p* < 0.05, \*\**p* < 0.01, \*\*\**p* < 0.001, \*\*\*\**p*<0.0001. **(C-H)** Western blot analysis of APP and p-tau levels in ReN cells treated with or without FA for 15 weeks after differentiation initiation. Tau phosphorylation levels were quantified, normalized to GAPDH, and the relative intensity in the control was set to 1. Results are presented as mean ± SEM (n = 3). \**p* < 0.05, \*\*\**p* < 0.001.

3D-cultured ReN cells, further supporting its potential role in AD initiation.

### Establishment of a 3D human neural cell culture AD model

Choi et al. first established a 3D human neural cell culture model of AD, which successfully recapitulates AD pathologies, including robust Aβ deposition and NFT-like tau pathology (Choi et al., 2014; Kim et al., 2015; Maniv et al., 2023). The 3D human neural cell culture AD model may more accurately mimic *in vivo* AD pathology than transgenic mouse models or 2D human cell cultures, providing a powerful platform for mechanistic studies of AD in a human-brain-like environment. Following the protocol established by Choi et al., we developed this model (Fig.4) to investigate the effects of FA on AD progression (Choi et al., 2014). Lentiviral particles expressing a GFP-APP containing both K670N/M671L (Swedish) and V717I (London) FAD mutations (APPSL) and an mCherry-PS1 carrying ΔE9 FAD mutation (PS1ΔE9) (AD model) were transduced into ReN cells in a 2D culture (Fig. 4). As a control, lentiviral particles expressing GFP and mCherry (G+M) were also transduced (Fig. 4).

**Figure 4.**
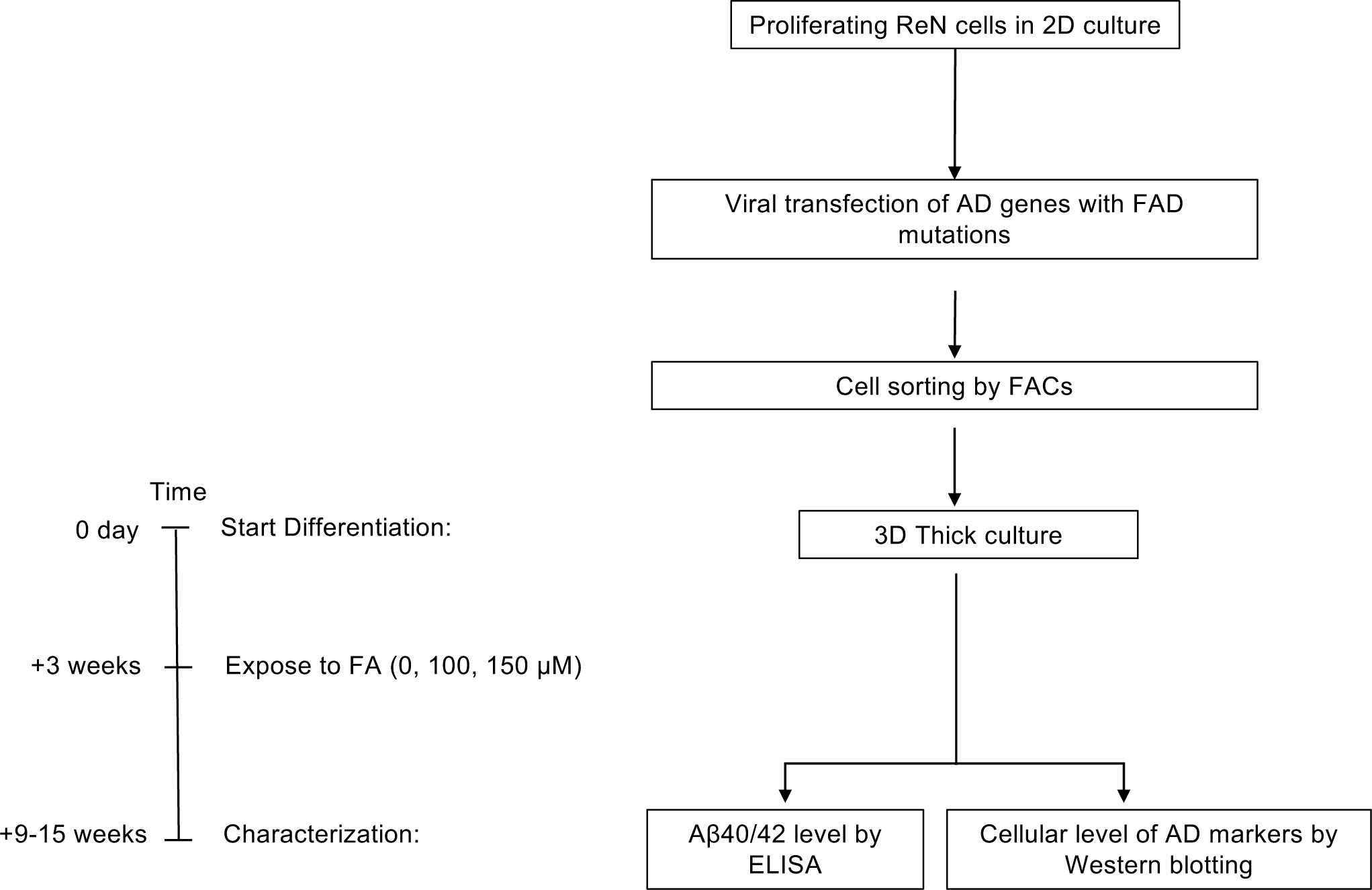
Flow chart for establishing a 3D-cultured human AD cell model.

Cells that highly expressed both exogenous proteins were collected by cell sorting, and their expression was confirmed by immunofluorescence (Fig. 5A). The sorted G+M and AD model cells were then plated onto the 3D Matrigel system and differentiated into neurons and glial cells by removing growth factors from the medium for three weeks (Fig. 4). We monitored AD markers for up to 12 weeks following differentiation. Aβ40 and Aβ42 levels were significantly increased in AD model cells compared to G+M control cells after 6 and 12 weeks of differentiation (Fig. 5B). Western blot analysis revealed that the levels of tau pathology markers, p-tau Ser199 and p-tau AT8, were dramatically elevated in AD model cells compared to G+M control cells 8 weeks after differentiation (Fig. 5C, D; Fig. S9). However, the increase in total tau levels was less pronounced (Fig. 5C, D; Fig. S9). Notably, the ratios of p-tau Ser199/total tau and p-tau AT8/total tau were significantly higher in AD model cells compared to the control, indicating that tau phosphorylation at different sites was increased (Fig. 5E). APP levels were also elevated in AD model cells compared to control cells (Fig. 5F, G; Fig. S10). These results confirm that the 3D human neural cell culture AD model was successfully established.

**Figure 5.**
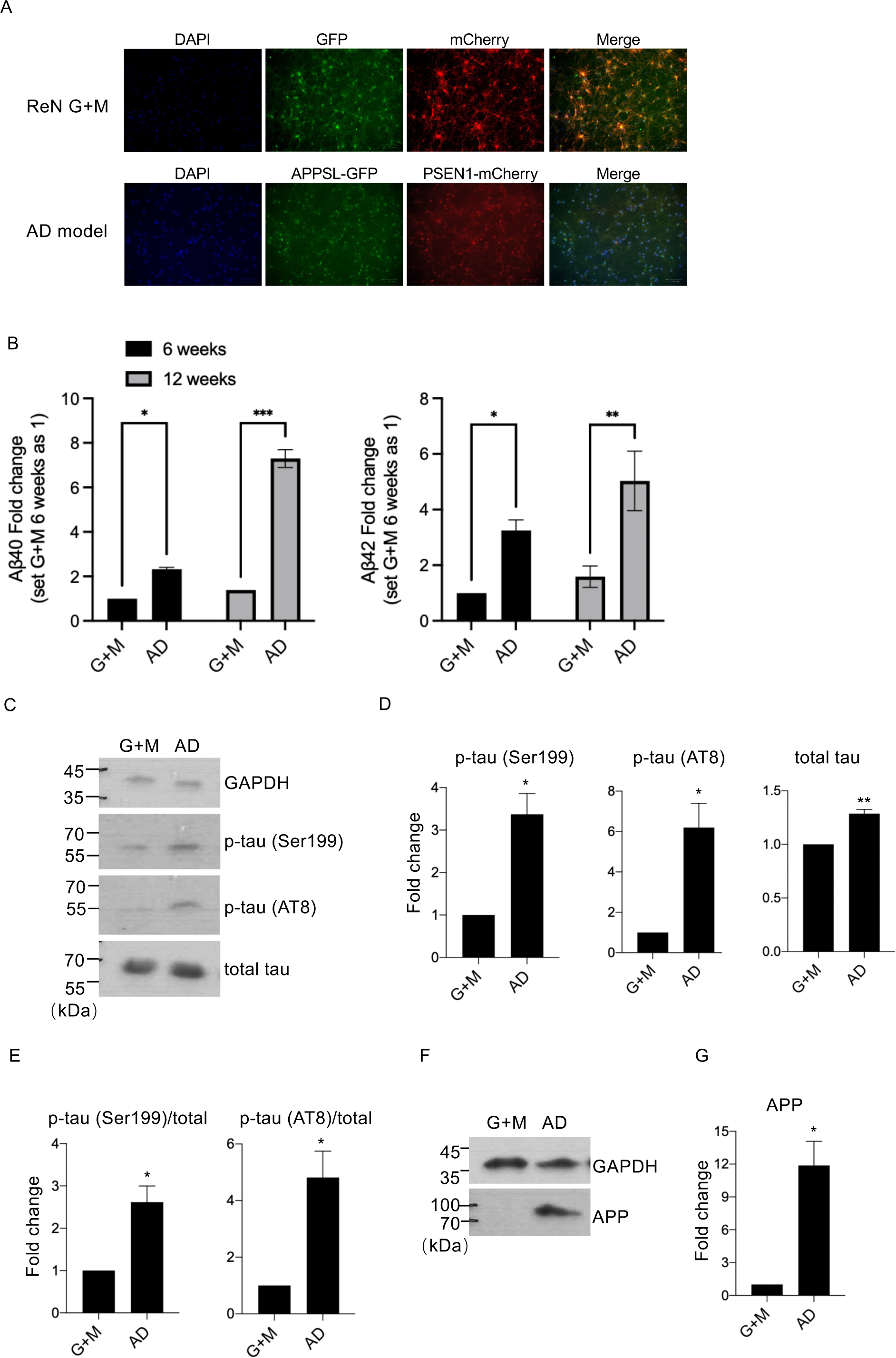
Establishment of a 3D-cultured AD model. **(A)** Immunofluorescence analysis of ReN cells transduced with lentiviruses expressing APPSL and PS1ΔE9, fused to fluorescent reporters. ReN cells were transduced with the indicated lentiviral constructs. Expression of APPSL-GFP (green) and PS1ΔE9-mCherry (red) was confirmed by immunofluorescence microscopy 8 weeks after differentiation initiation. **(B)** Analysis of soluble Aβ40 and Aβ42 levels. The increased Aβ40 and 42 levels were measured by ELISA at different timepoints (6 and 12 weeks) and normalized to protein concentration. The relative level of Aβ40 and Aβ42 in G+M at 6 weeks was set to one. Unpaired t test was used to compare differences between G+M and AD models at different time points. \**p* < 0.05, \*\**p* < 0.01, \*\*\**p* < 0.001. **(C–G)** Western blot analysis of tau phosphorylation at different sites, total tau, and APP levels. Eight weeks after differentiation initiation, 3D-cultured G+M and AD model cells were collected, and whole cell lysates were extracted for Western blot analysis (C and F). The levels of p-tau, total tau, and APP were quantified, normalized to GAPDH, and the relative intensity in G+M was set to one (D, G). Tau phosphorylation levels at two different sites normalized to total tau, to calculate the p-tau vs. total tau ratio (E). Unpaired t test was used to compare differences between G+M and AD models at different time points. Results are presented as mean ± SEM (n = 3). \**p* <0.05 and \*\**p* < 0.01.

### FA exposure facilitates AD progression in the human neural cell 3D AD model

To investigate the role of FA in AD progression, 0, 100, and 150 μM FA were added to the culture medium of G+M control and AD model cells three weeks after differentiation initiation. The levels of Aβ40 and Aβ42 were monitored using ELISA after 6 and 12 weeks of FA exposure. Aβ40 levels were significantly increased in AD model cells following exposure to 100 and 150 μM FA at both 6 and 12 weeks (Fig. 6A, B). Similarly, Aβ42 levels were elevated in a dose-dependent manner upon FA exposure at both time points (Fig. 6C, D). Western blot analysis further revealed that p-tau Ser199 levels were dose-dependently increased following FA exposure, and tau phosphorylation at Ser202&Thr205 was significantly increased upon exposure to 150 μM FA (Fig. 6E-H; Figs. S11 and S12). Additionally, the potential AD pathology marker APP was also found elevated in response to FA treatment (Fig. 6I, J; Fig. S13). These results suggest that FA exposure may accelerate AD progression in the human neural cell 3D AD model.

**Figure 6.**
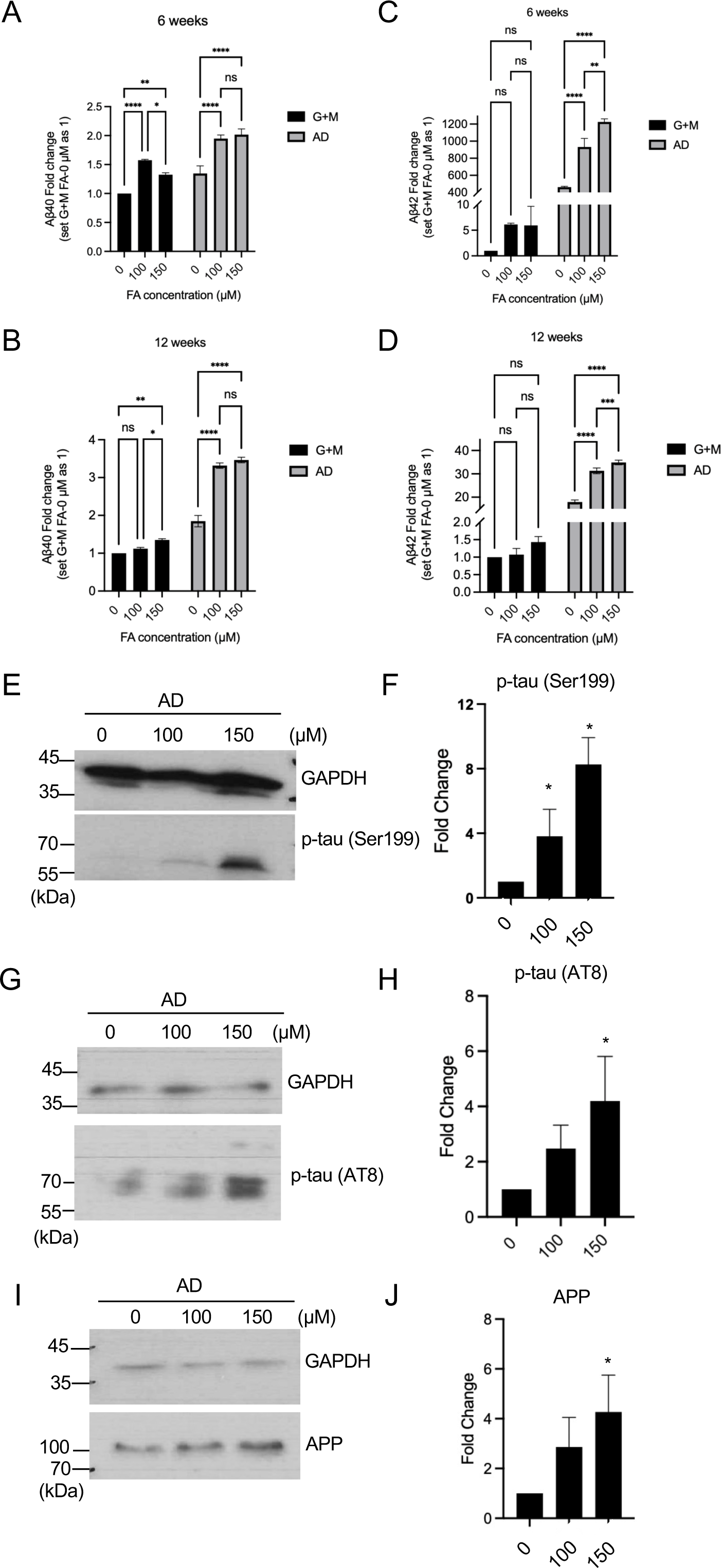
Analysis of soluble Aβ levels in the 3D-cultured AD models exposed to FA. After differentiation, G+M and AD model cells were treated with FA (0, 100, and 150 μM) for 6 or 12 weeks. **(A, B)** Aβ40 levels in the medium were measured by ELISA at 6 and12 weeks. **(C, D)** Aβ42 levels in the medium were measured by ELISA at 6 and 12 weeks. Two-way ANOVA suggested significant interaction between Aβ concentration and G+M or AD. Turkey’s multiple comparisons test was used to analyze the effect of FA on G+M and AD groups. \**p* < 0.05, \*\**p* < 0.01, \*\*\**p* < 0.001, \*\*\*\**p* < 0.0001. **(E-J)** Western blot analysis of APP and p-tau levels in AD model cells with or without FA exposure. One-way ANOVA, followed by Turkey’s multiple comparisons test, was used to analyze the effect of FA on G+M and AD groups. Results are presented as mean ± SEM (n = 3). \**p* < 0.05.

## Discussion

We investigated the role of FA exposure in AD pathologies using a human neural cell 3D culture system and a human neural cell 3D AD model, demonstrating that FA exposure can induce and accelerate both Aβ and tau pathologies in human neural cells. This is the first study to highlight the significance of FA in AD development using 3D-cultured human neural cells.

With the ability to penetrate the blood-brain barrier (BBB), both exogenous and endogenous FA can accumulate in the brain, potentially leading to brain disorders. Exogenous FA has been correlated with cognitive impairment in occupationally exposed cohorts (Kilburn et al., 1987; Kilburn & Warshaw, 1992; Letellier et al., 2022; Zendehdel et al., 2016) as well as in animal models exposed to FA (Liu et al., 2018; Tong, Han, Luo, Wang, et al., 2013; Zhai et al., 2018). Endogenous formaldehyde levels in the brain and urine have been found to increase with aging and accumulate in patients with age-related dementia (Kou et al., 2022) and AD (Tong et al., 2017; Tong et al., 2011). Moreover, endogenous FA levels are positively correlated with the severity of cognitive impairments (Kou et al., 2022). These studies suggest that excessive FA accumulation in the brain, whether endogenous or exogenous, plays a critical role in cognitive dysfunction and neurotoxicity.

FA has been linked to various AD pathologies, particularly its interaction with Aβ peptides. Studies have shown that FA and Aβ mutually influence each other. Excessive FA exposure can induce Aβ aggregation in the brains of primates and rodents (Liu et al., 2018; Zhai et al., 2018). Additionally, FA can directly increase the size of freshly prepared seed-free Αβ_1−40_ peptides (Chen et al., 2006). In turn, Aβ contributes to endogenous FA accumulation by inactivating FA dehydrogenase (FDH) (Yue et al., 2019). In our study, APP protein, the precursor of Aβ peptides, was significantly elevated following FA exposure in both 2D- and 3D-cultured differentiated ReN cells. Furthermore, Aβ40 and 42 levels were dramatically increased in 3D-cultured ReN cells and AD model cells in a dose-dependent manner. These findings suggest that excessive FA not only triggers Aβ aggregation, potentially initiating AD onset, but also accelerates the Aβ cascade during the AD progression.

FA has been shown to induce tau misfolding and aggregation, leading to formation of tau aggregates that display neurotoxicity (Nie et al., 2007). Later study found that FA exposure causes tau hyperphosphorylation (in both cytoplasmic and nuclear fraction) and promotes tau polymerization in neuroblastoma (N2a) cells and mouse brains (He et al., 2016; Lu et al., 2013). In our study, one-week exposure to 100 μM FA increased tau phosphorylation at multiple sites, including Ser199, Ser202, and Thr205, without changing the total tau levels in 2D-cultured differentiated ReN cells. An increase in tau phosphorylation at Ser199 was also observed in rat brain tissues following nasal exposure to 2 ppm FA for 28 days. Furthermore, in 3D-cultured differentiated ReN cells and AD model cells, p-tau levels increased dramatically in a dose dependent manner following FA exposure. These findings demonstrate that FA induces and accelerates AD-related tau pathologies both *in vitro* and *in vivo*.

The relationship between Aβ and tau pathologies has been a topic of debate for decades but it is likely that these pathologies have synergistic effects, which fundamentally drive disease progression (Busche & Hyman, 2020). In our study, we observed the co-presence of both Aβ and tau pathologies in 3D-cultured AD model cells carrying APP/PSEN1 mutations, including an increase in APP and Aβ peptides, as well as increased total tau levels and tau hyperphosphorylation at multiple sites. Moreover, FA exposure simultaneously induced Aβ and tau pathologies in both 2D- and 3D-cultured normal neural cells, while accelerating these pathologies in 3D-cultured AD model cells. These findings suggest that emerging therapeutic strategies targeting upstream events common to Aβ and tau pathologies–or disrupting their pathological synergies–may be more effective than approaches that target them independently. Furthermore, eliminating FA as a trigger or an accelerator of AD pathologies, or targeting key mediator in FA-mediated AD progression, could provide novel therapeutic avenues for approaches for AD treatment.

This study demonstrated that FA induces AD-like pathologies not only in human neural cells but also in rat brains. The ability of ReN cells to differentiate into neurons and glial cells, including astrocytes and oligodendrocytes, makes them a closer approximation to multicellular brain environment compared to monocellular culture. The 3D-cultured system provides ReN cells with a solid environmental that support growth and cell-cell interactions. Additionally, it retains secreted Aβ peptides, maintaining high local concentration, which are easily lost during medium changes in 2D cultures. In our study, we monitored markers for both Aβ pathologies and tau pathologies, as those are closely linked in AD. We found that FA induces both pathologies in normal differentiated human ReN cells, suggesting potential adverse effects of FA exposure in healthy individuals._However, our ReN cell culture model lacks microglial cells, which play a crucial role in Aβ clearance. Furthermore, as Aβ can induce elevated levels of FA, this system does not distinguish between endogenous and exogenous FA.

## Conclusion

In this study, we demonstrate that FA exposure induces AD-like pathologies in differentiated human neuron progenitor cells as well as in rat brains. FA exposure contributes to both initiation and acceleration of AD pathology cascades in 3D-cultured ReN cells and AD model cells, respectively. These findings suggest that FA toxicity is closely linked to key hallmarks of AD pathology and plays a significant role in AD pathogenesis. Further investigation into the underlying mechanisms may provide valuable insights for AD prevention and therapeutic strategies.

## Acknowledgement

This work was supported by a grant from the US National Institutes of Health (R01ES030583 to C.J.).

