## Supplemental Figures for "Formaldehyde induces and promotes Alzheimer’s disease pathologies in a 3D human neural cell culture system"

### Figure S1: Full-length Western blot images related to Figure 1B.

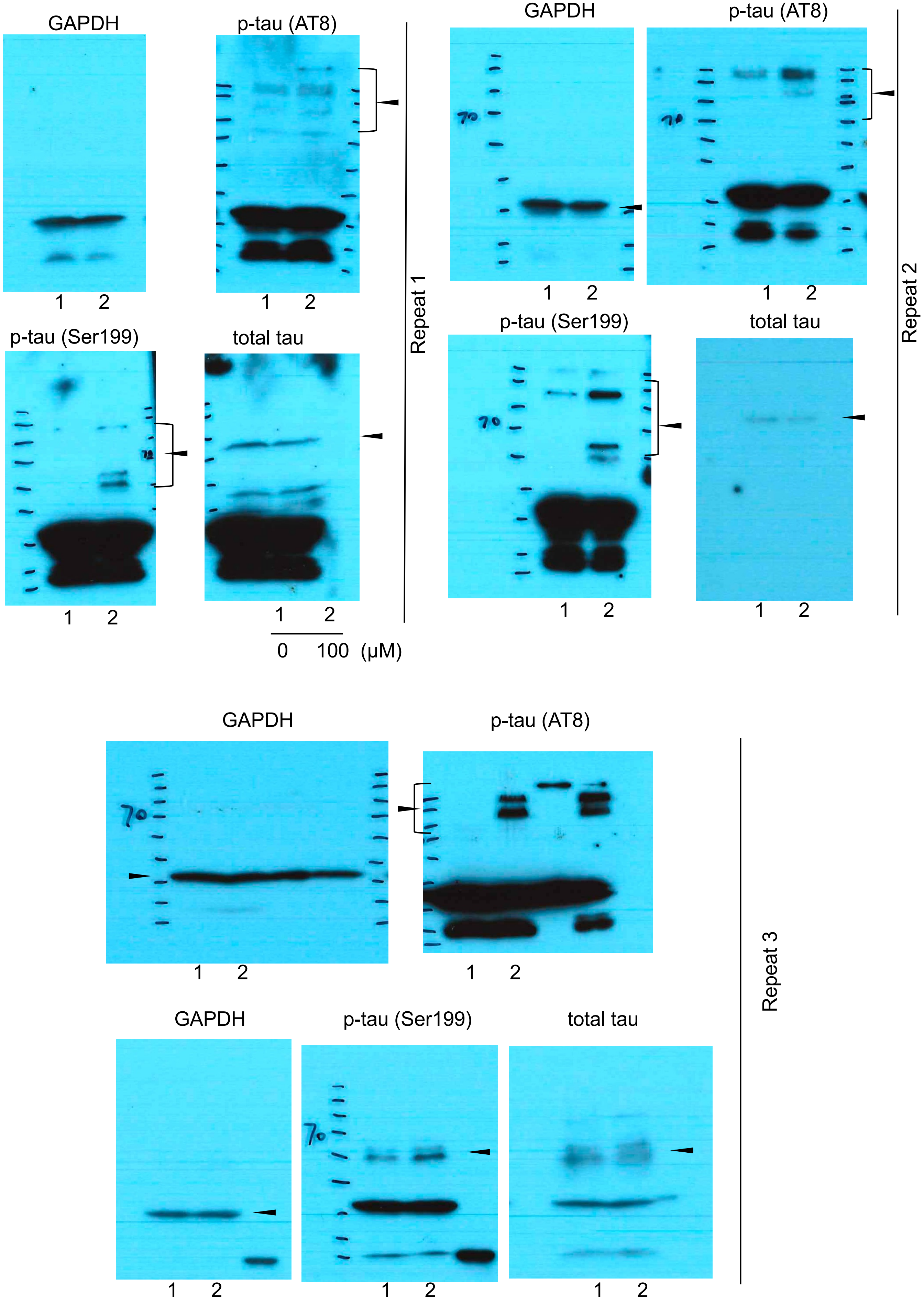

Figure S2: Full-length Western blot images related to Figure 1D.

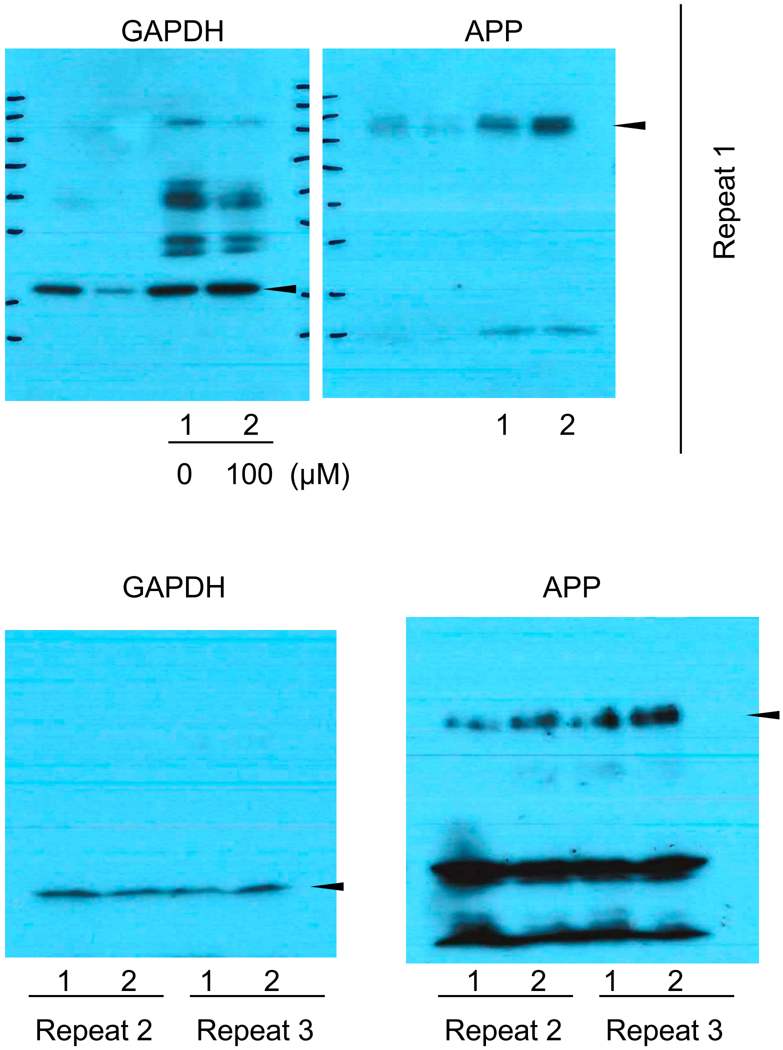

Figure S3: Full-length Western blot images related to Figure 2A.

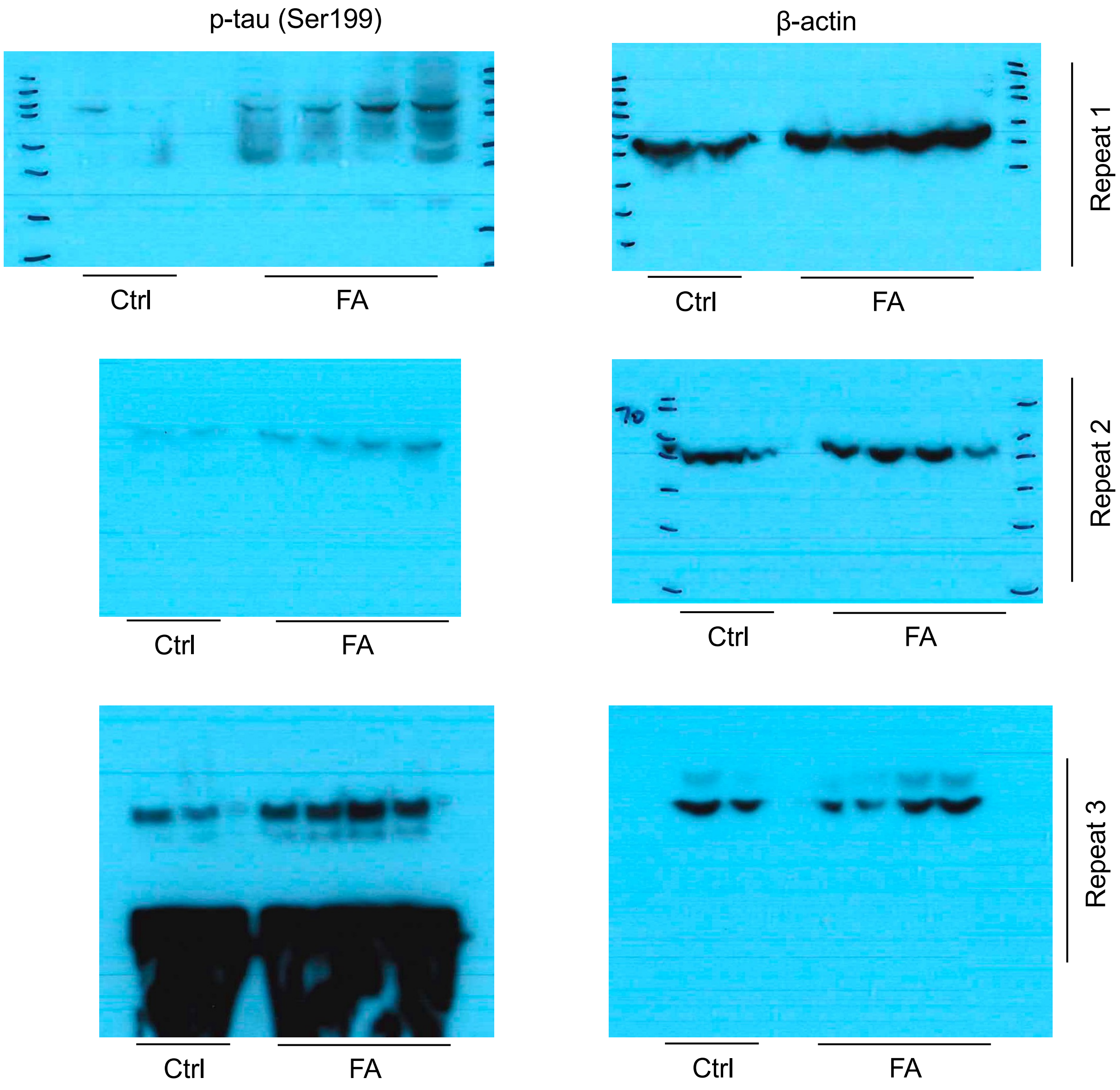

Figure S4: Full-length Western blot images related to Figure 2C.

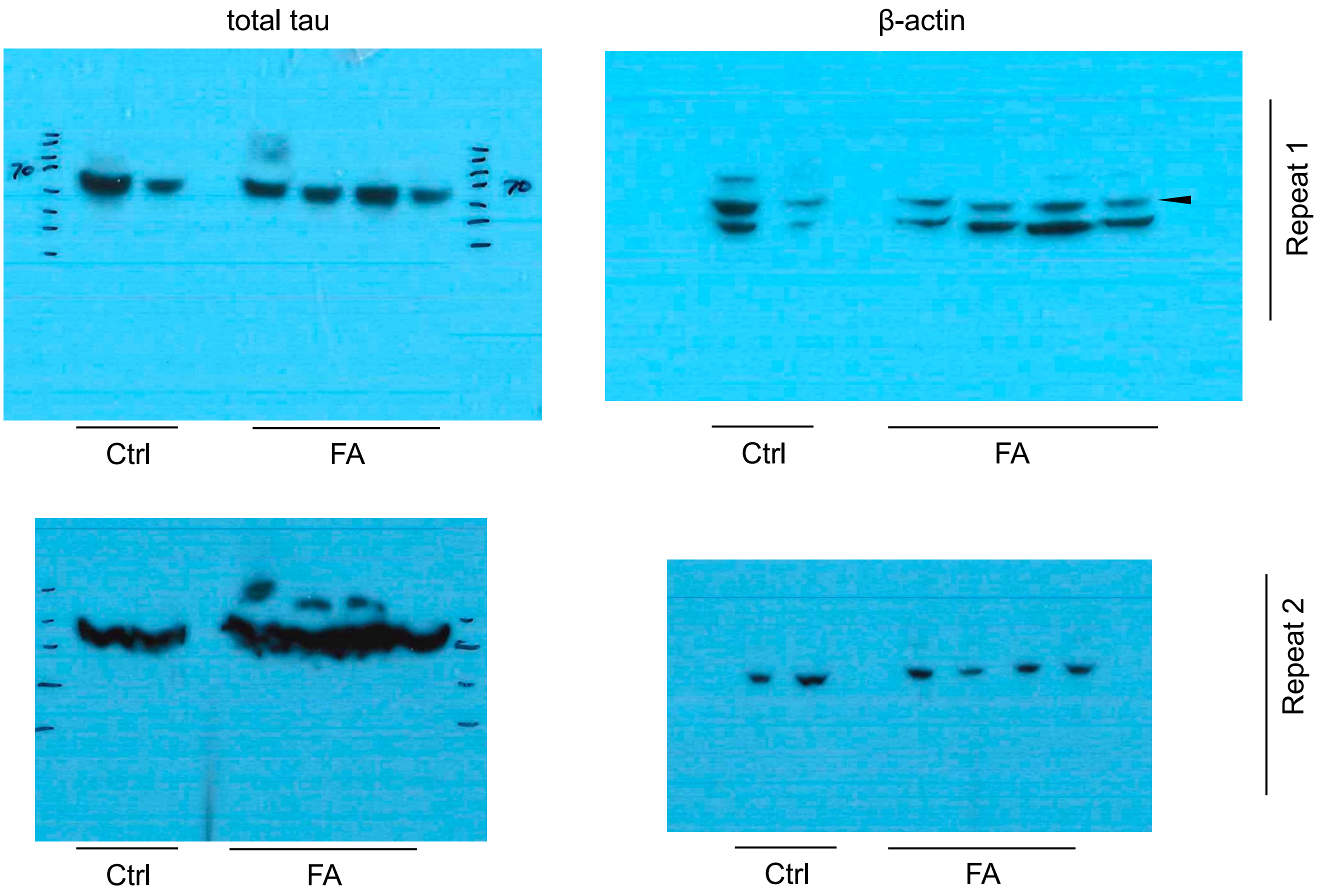

Figure S5: Full-length Western blot images related to Figure 2E.

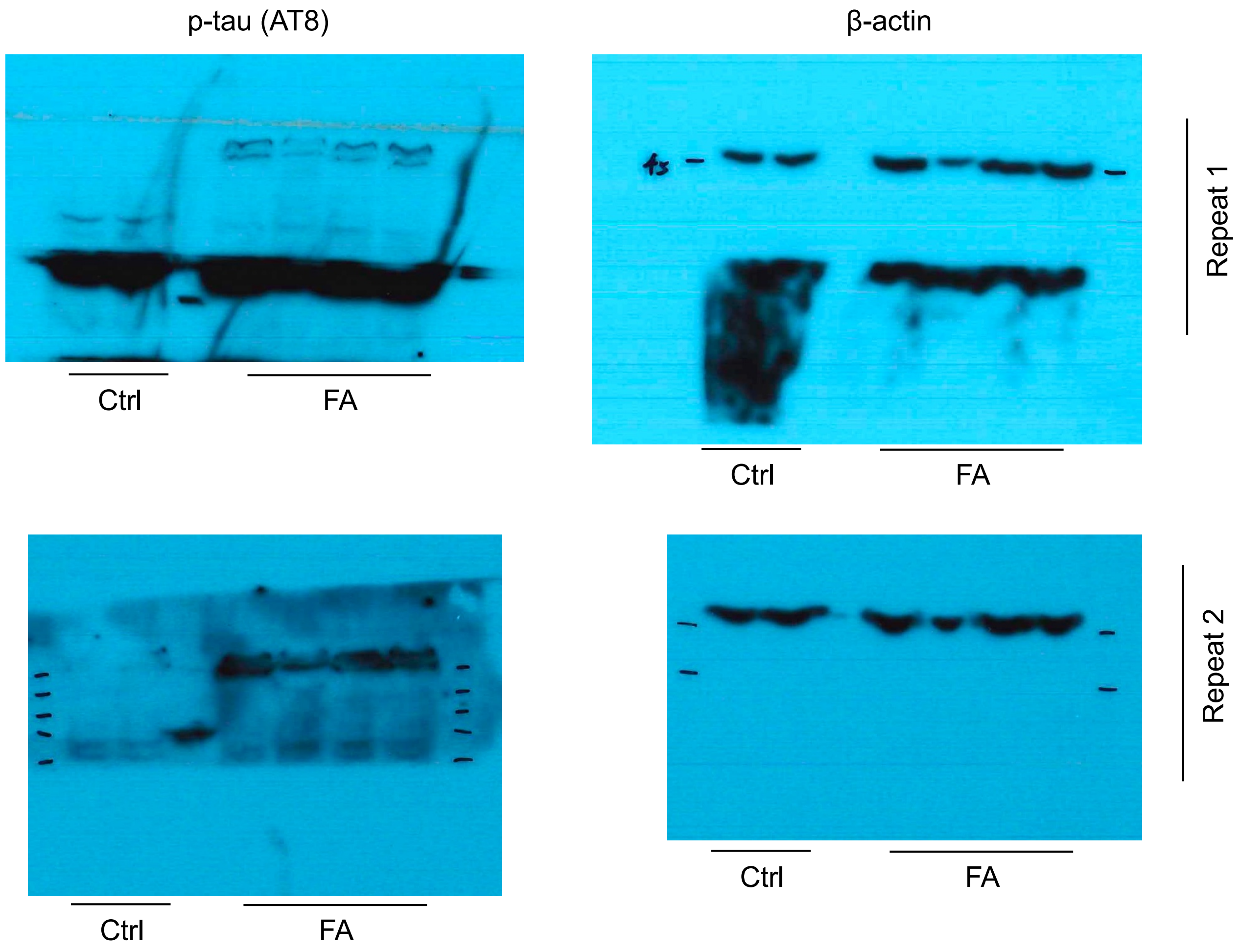

Figure S6: Full-length Western blot images related to Figure 3C.

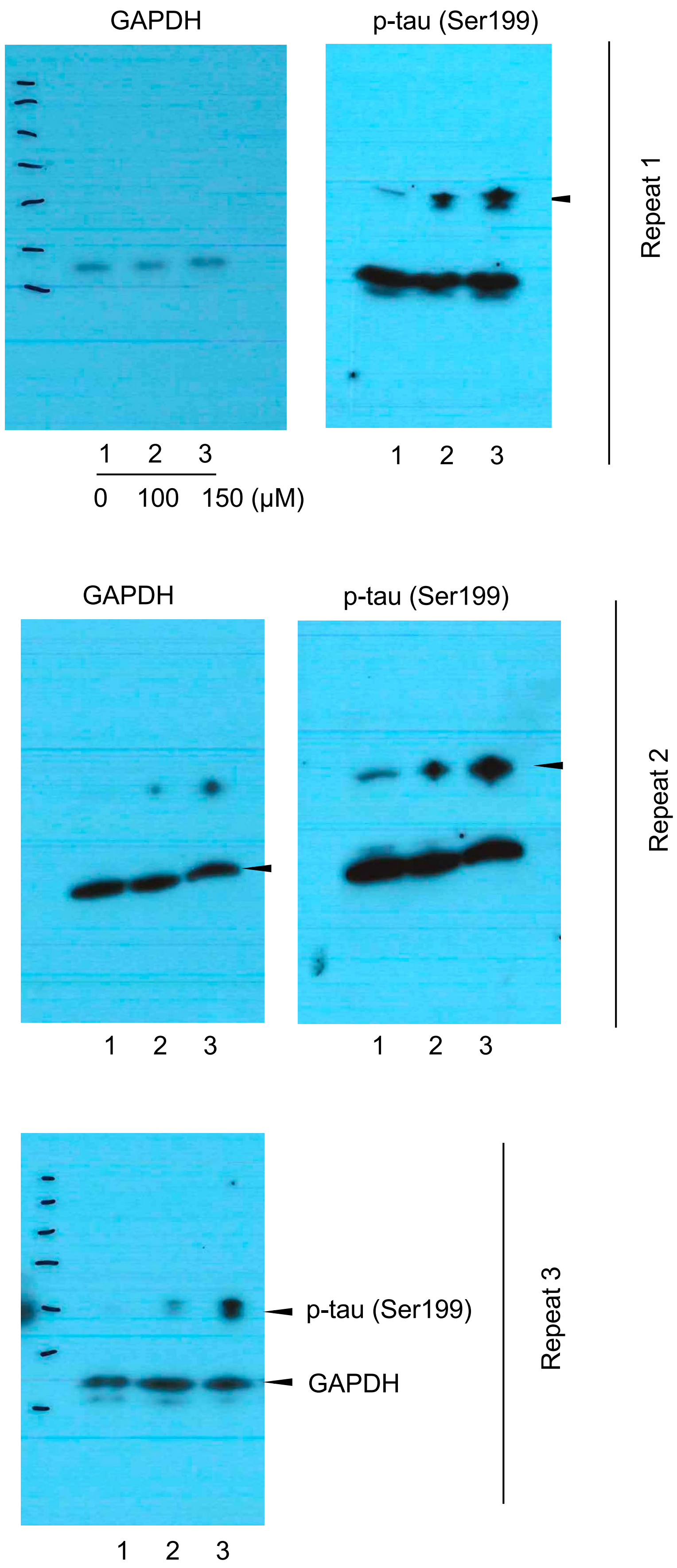

Figure S7: Full-length Western blot images related to Figure 3E.

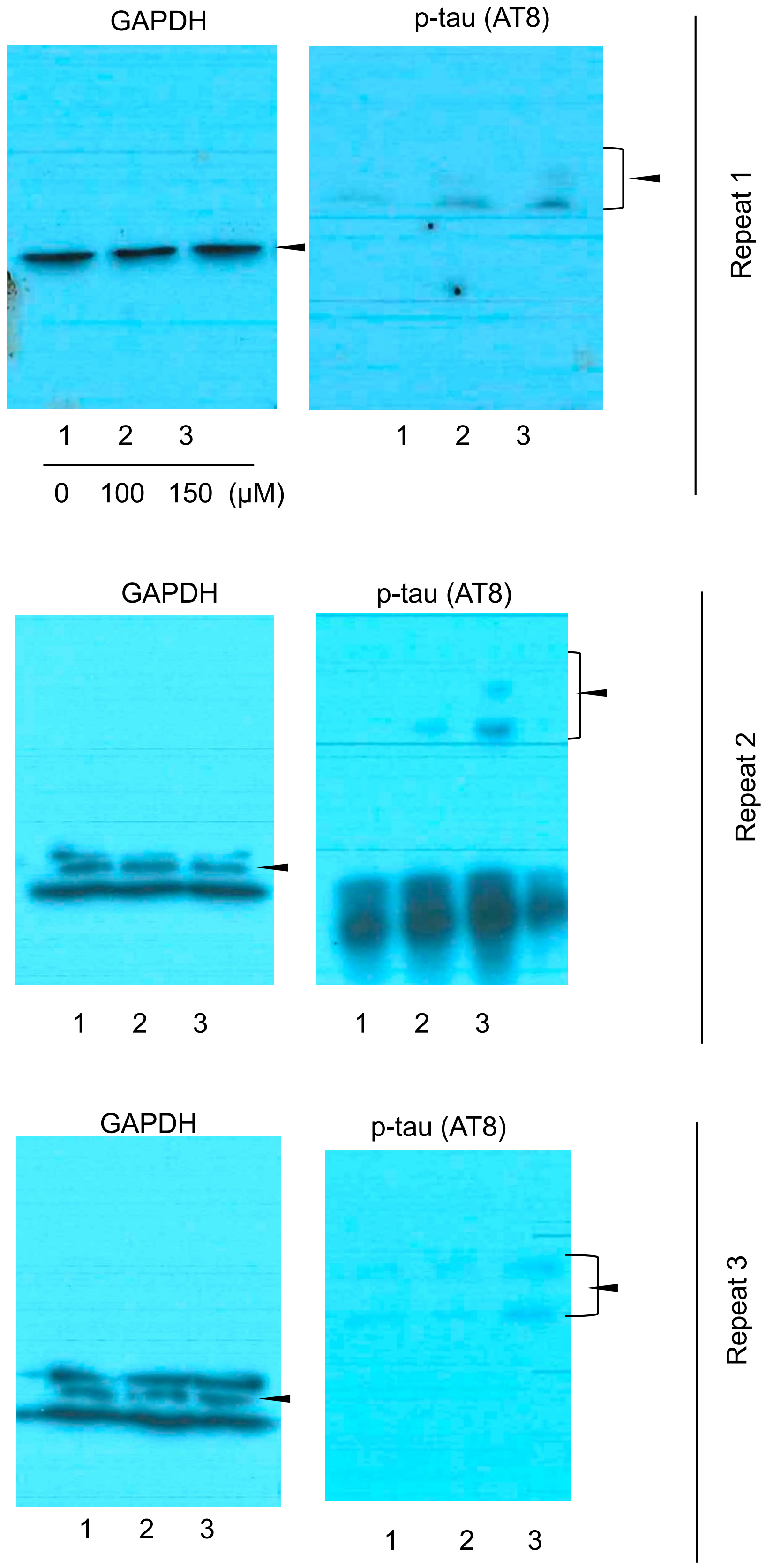

Figure S8: Full-length Western blot images related to Figure 3G.

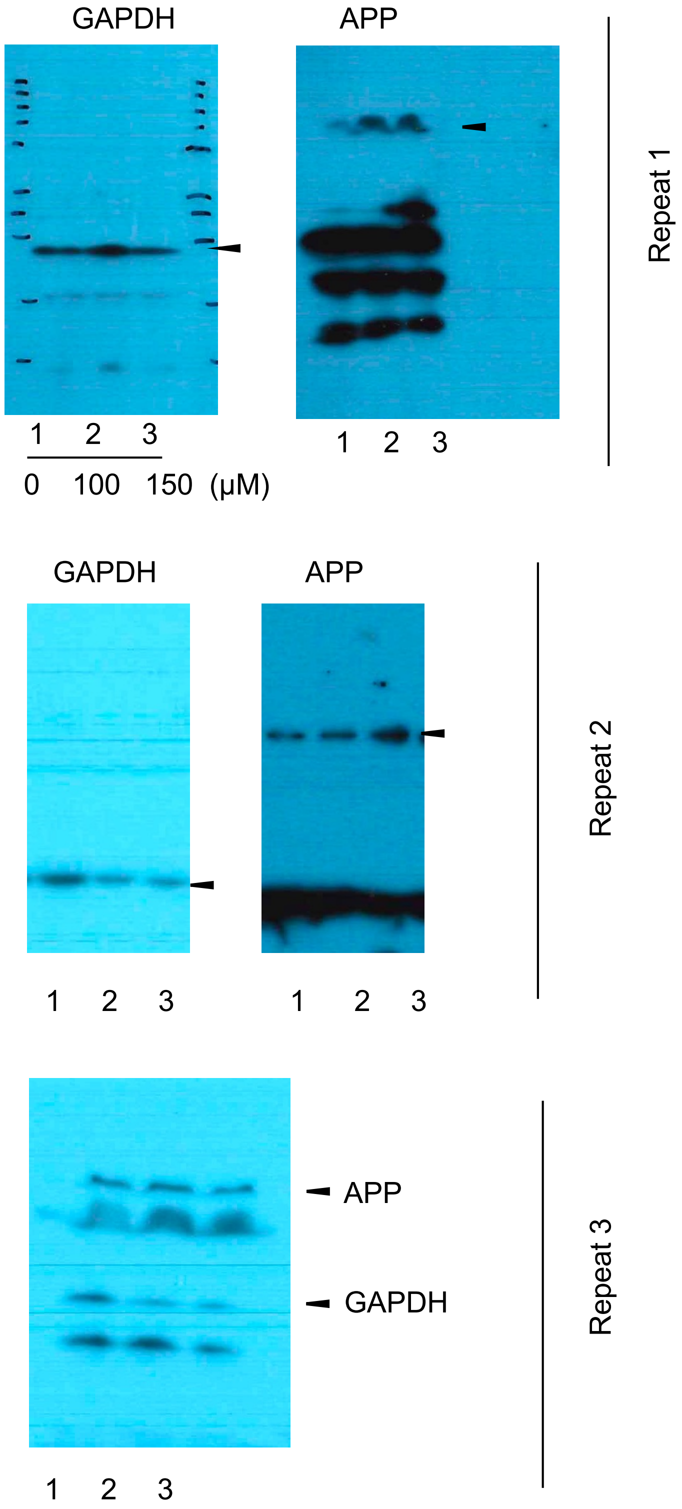

Figure S9: Full-length Western blot images related to Figure 5C.

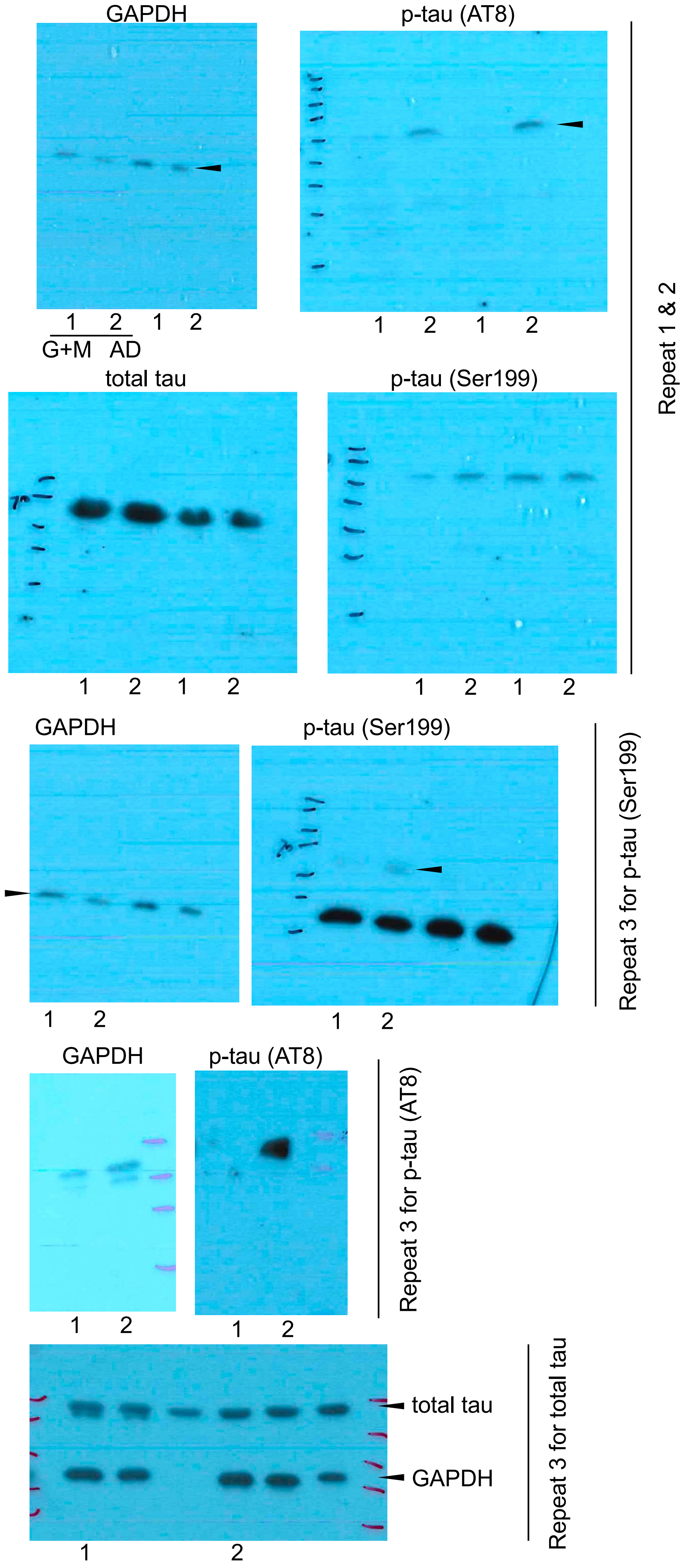

Figure S10: Full-length Western blot images related to Figure 5F.

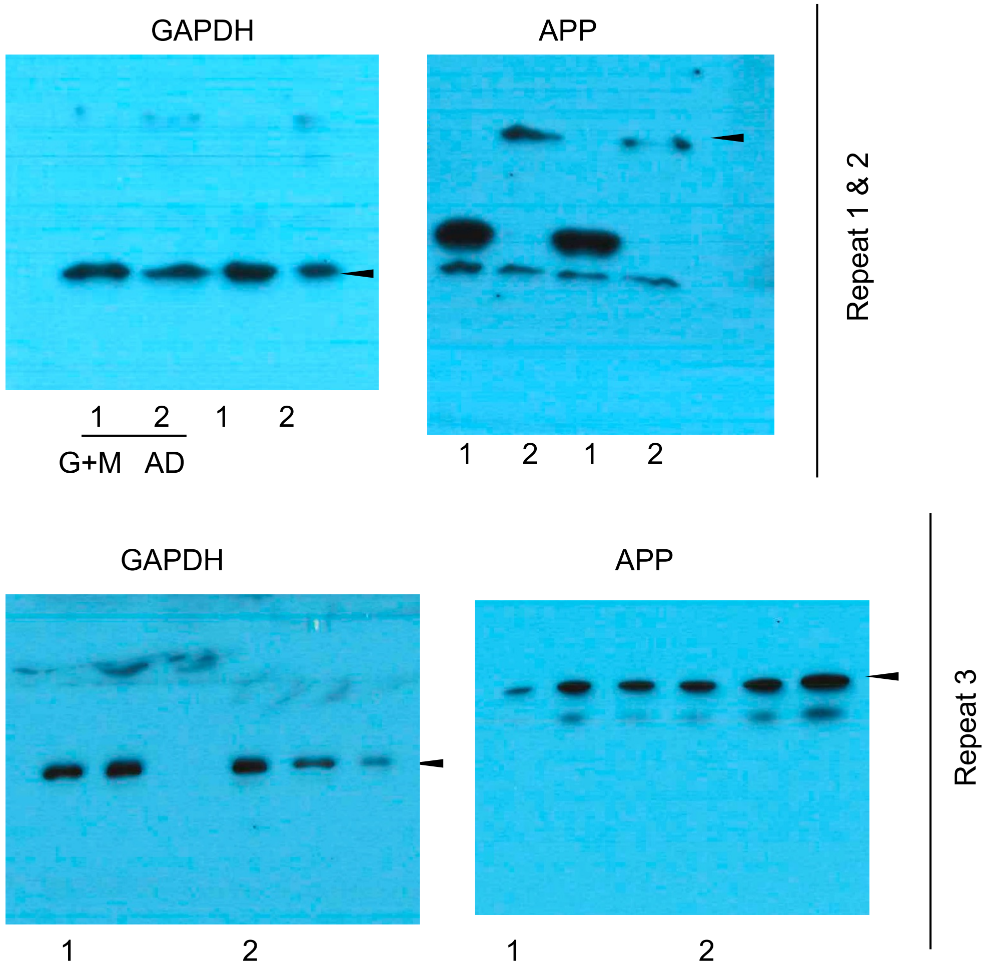

Figure S11: Full-length Western blot images related to Figure 6E.

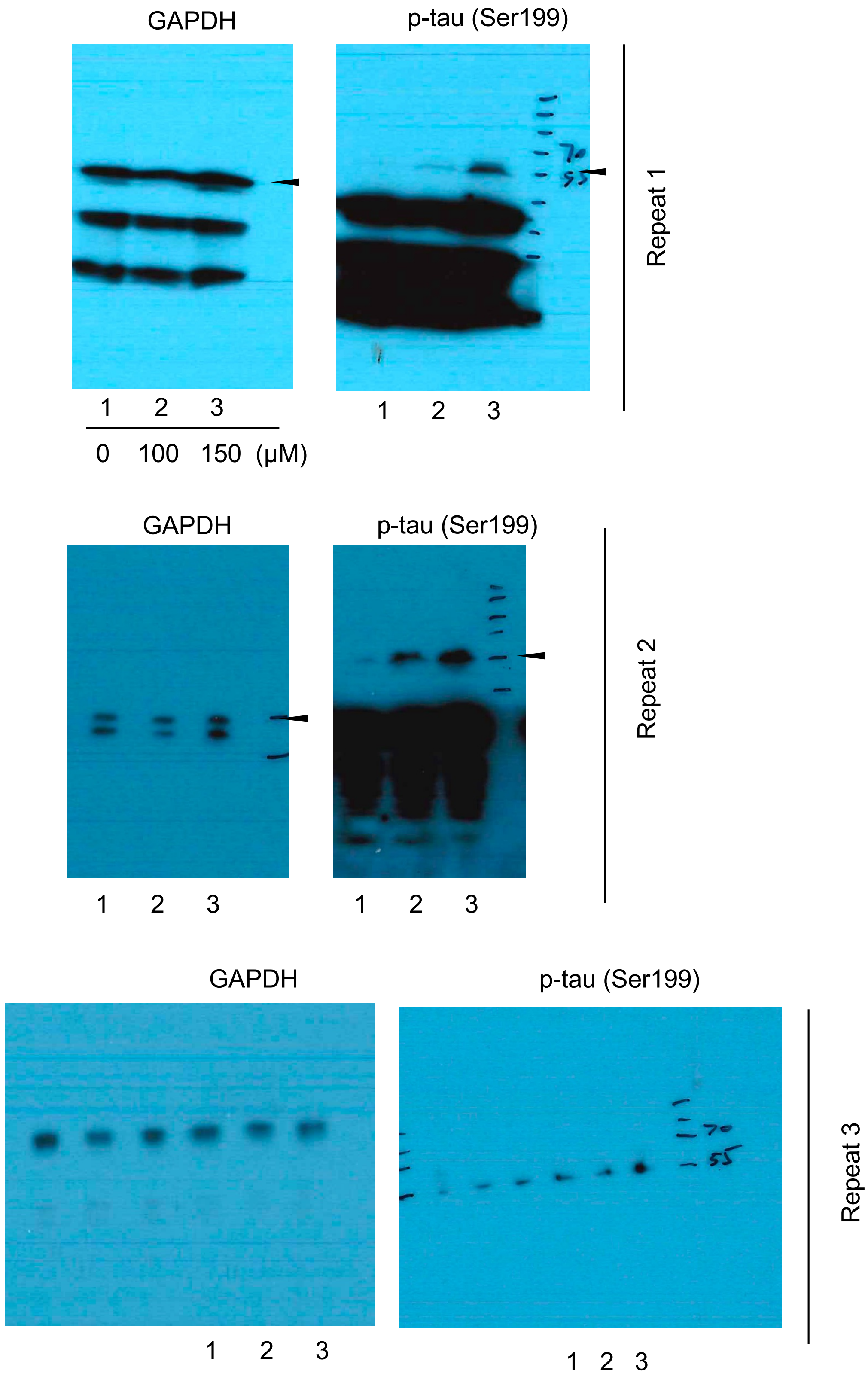

Figure S12: Full-length Western blot images related to Figure 6G.

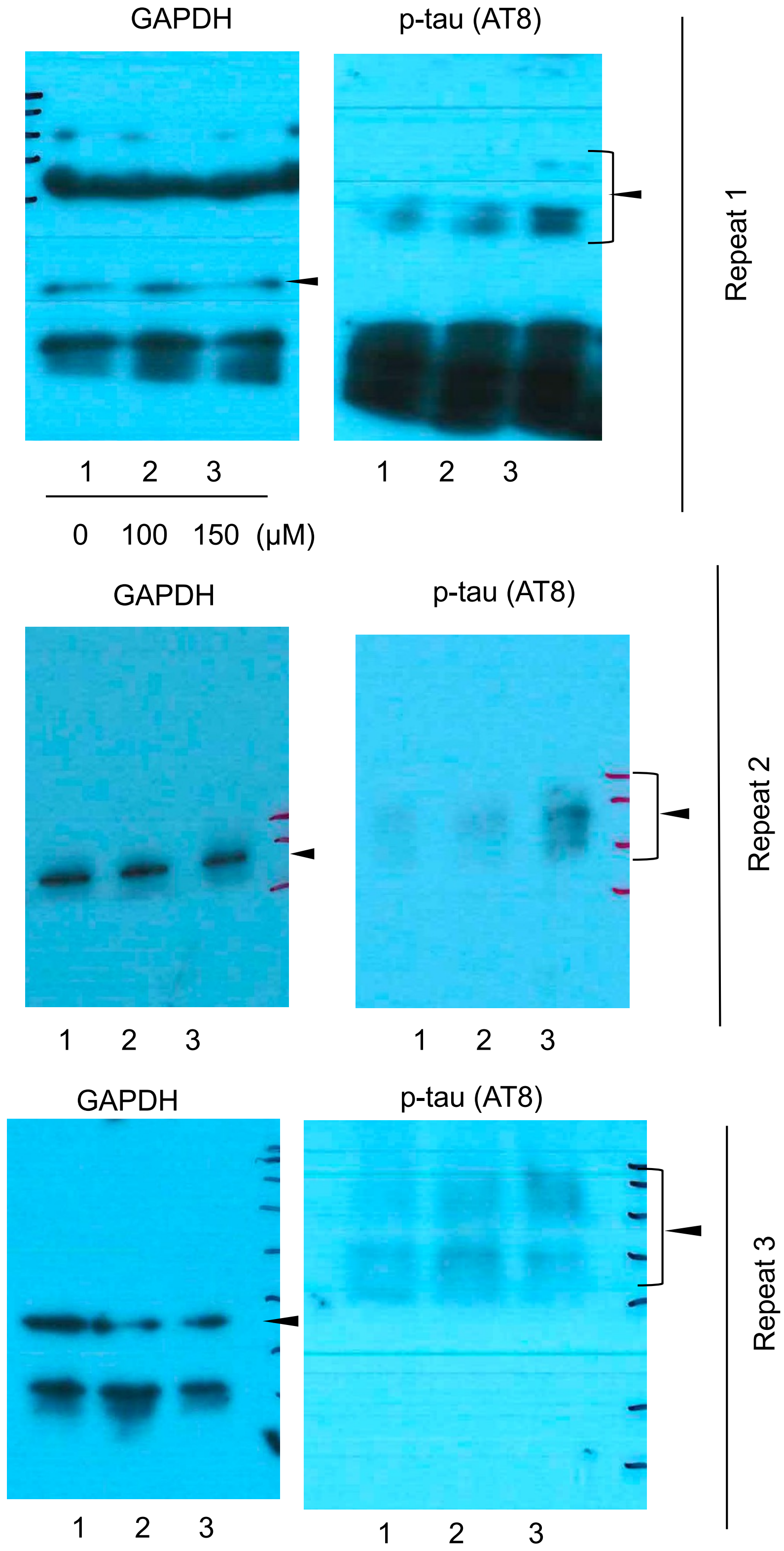

Figure S13: Full-length Western blot images related to Figure 6I.

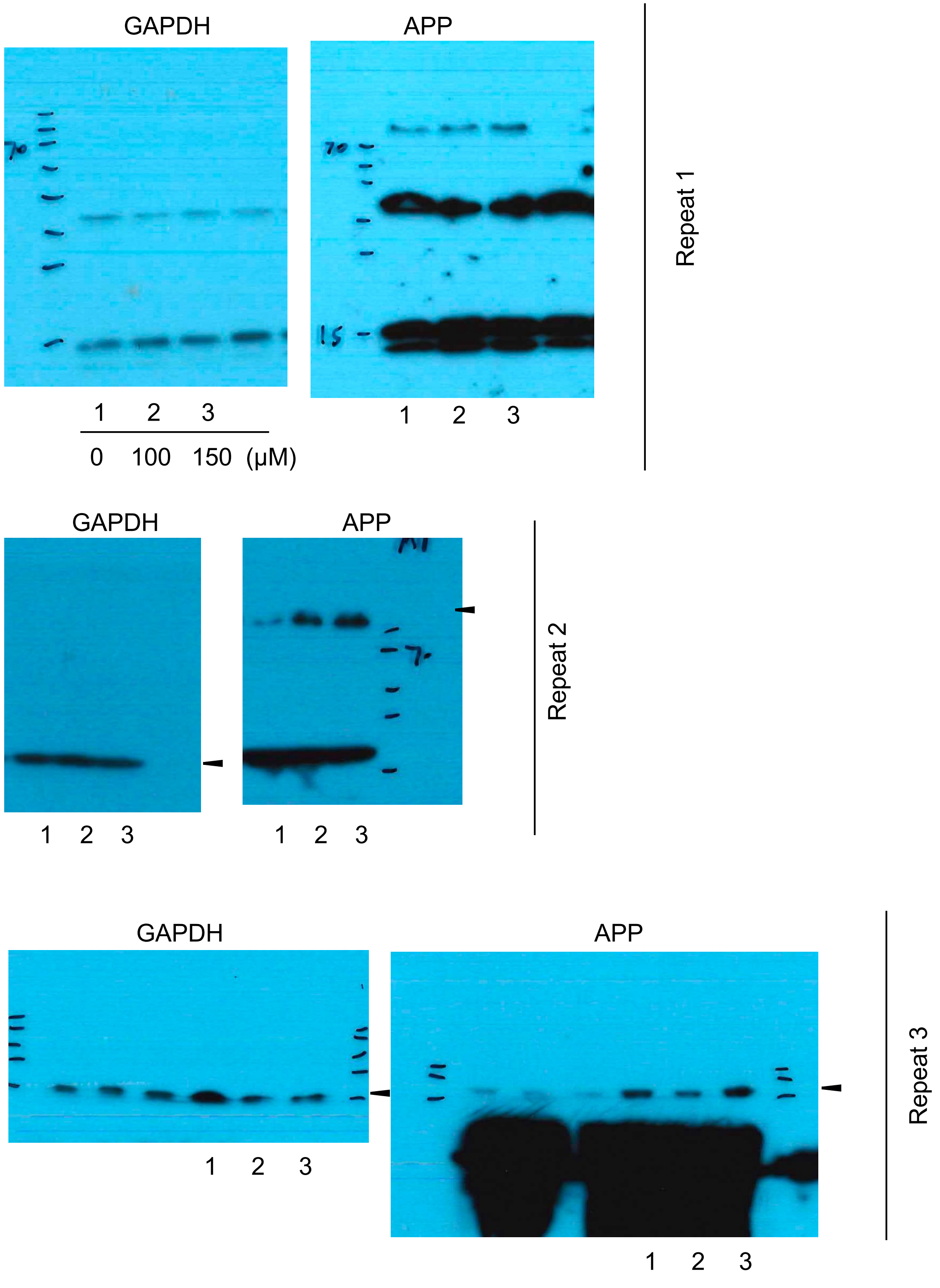
